# Molecular characterization of baculovirus-induced chromatin marginalization and architectural alteration

**DOI:** 10.1101/2023.07.17.549271

**Authors:** Xiangshuo Kong, Guanping Chen, Conghui Li, Xiaofeng Wu

**Affiliations:** College of Animal Sciences, Zhejiang University, Hangzhou 310058, China; Key laboratory of silkworm and bee resource utilization and innovation of Zhejiang Province, Hangzhou, China

**Keywords:** chromatin marginalization, Lamin A/C, HP1a, nuclear actin, genome structure, baculovirus

## Abstract

To facilitate rapid replication and assembly of progeny, baculovirus is known to manipulate the host nuclear microenvironment by inducing chromatin changes in localization and architecture. However, the molecular mechanisms underlying these changes remain unknown. Here, we revealed that the nuclear lamina (NL) protein Lamin A/C interacts with the heterochromatin protein 1 alpha (HP1a) and identified the middle region of HP1a as critical for this interaction. Suppression of Lamin A/C and HP1a expression resulted in a significant inhibition of chromatin marginalization mediated by baculovirus infection. Moreover, the heterochromatin modification H3K9me3, which is recognized and bound by HP1a, also participated in the process of chromatin marginalization. Our live-cell imaging and quantitative analysis unveiled a passive function of marginal chromatin, which involves the formation of a physical barrier that impedes the nuclear egress of the nucleocapsids. Furthermore, baculovirus-induced nuclear F-actin altered the steady-state of intranuclear actin pool, thus regulating the nucleosome disassembly. Overall, our findings illustrate the molecular mechanisms dictating chromatin marginalization and structural alterations during baculovirus infection, shedding new light on the potential function of marginalized chromatin in the origin of its biphasic life cycle.

**Author Summary:** In our previous study, we illustrated the organization and accessibility of chromatin marginalized by baculovirus infection through a combination of ATAC-seq and biochemical assays. Here, we further dissect the molecular mechanism underlying the baculovirus infection induced chromatin marginalization and disassembly. Specifically, baculovirus utilizes the Lamin A/C-HP1a-H3K9me3 axis to mediate chromatin marginalization at the nuclear periphery. When the interaction between Lamin A/C and HP1a is disrupted, the marginalization of chromatin is also affected. Furthermore, our single-virion tracking results indicate that the marginalized chromatin forms a physical barrier, impeding the nuclear export of nucleocapsids at the very late stage of infection. For the changes in chromatin architecture, we propose a model in which baculovirus infection induced nuclear F-actin compromises the dynamics of nuclear actin pool, which in turn promotes chromatin disassembly. Taken together, we provide a comprehensive understanding of molecular mechanism of baculovirus infection induced changes in chromatin localization and organization, which lay the foundation for studies on how DNA viruses manipulate the nuclear microenvironment.

## Introduction

Typically, DNA viruses replicate and assemble in the host nucleus, which inevitably alters the spatial and structural organization of nucleus. A prominent hallmark of DNA virus infection is the formation and expansion of viral replication compartment (VRC) in the nucleus^1^, which is accompanied by changes in the host genome including chromatin remodeling and re-localization^2–5^. These alterations are highly conserved from invertebrate to vertebrate DNA viruses. As the most well-studied representatives of invertebrate and vertebrate DNA viruses, baculovirus and herpes simplex virus 1 (HSV-1), respectively, both induce chromatin remodeling and re-localization (marginalization) during infection. Baculovirus is a diverse group of invertebrate-specific viruses with double-stranded DNA genome^6^. Similar to other DNA viruses, baculovirus infection results in the formation of virogenic stroma (VS), the VRC of baculovirus, within the host nucleus, and also causes significant chromatin marginalization^7^. During its life cycle, it produces two different virions, budded virion (BV) and occlusion-derived virion (ODV). BV is responsible for the cell-to-cell infection within an organism, while ODV is responsible for the oral infection to initiate the primary infection^8^. Currently, there is still no consensus on the origin of this unique biphasic life cycle, in other words, it is a chicken-and-egg problem.

Recently, baculovirus infection induced changes in the accessibility and organization of marginalized chromatin have been characterized^4^. The data suggested that the marginalized chromatin genome-widely gains accessibility with disassembly of multi-nucleosomes. The potential roles of host nuclear lamina and heterochromatin protein 1 alpha (HP1a, also known as CBX5) in mediating the tethering of host chromatin to the nuclear periphery were discussed. NL is a dense filamentous meshwork underlying the nuclear membrane principally composed by the Lamin proteins^9^. In addition to functioning as the nucleoskeleton, Lamin proteins can also serve as a scaffold for orderly organizing the heterochromatin at the nuclear periphery through interacting with Lamin B receptor (LBR) and HP1a^10, 11^. In mammals, the Lamin proteins are grouped into two classes: A-type (Lamin A/C) and B-type (Lamin B1, B2 and B3)^12^. Within the host of baculovirus, most invertebrates only possess a single *Lamin* gene in their genomes. Such as, the domestic *Bombyx mori* genome contains a *Lamin A/C* gene but lacks the B-type Lamin proteins^13^, which will provide a simple genetic background to reveal the function of Lamin protein in baculovirus infection induced chromatin marginalization. Hence, we utilize the Bombyx mori nucleopolyhedrovirus (BmNPV), a representative member of baculovirus, as a model to investigate the molecular mechanism of DNA virus infection induced chromatin marginalization.

In the past decade, it has become clear that the nuclear actin plays an essential part in many fundamental nuclear processes, including transcription regulation and initiation^14^, chromatin reorganization and remodeling^15, 16^, DNA damage repair^17, 18^, and maintenance of nuclear architecture^19^. Specifically, a recent article identified a cell-cycle controlled nuclear F-actin that reorganizes the mammalian nucleus and chromatin assembly after mitosis^20^. Moreover, human cytomegalovirus (HCMV) infection induced nuclear F-actin causes extensive spatial and organizational polarization of host chromatin within the nucleus^21^. These findings remind us of a hallmark of baculovirus infection, in which the remarkable formation of nuclear F-actin mediates the nuclear egress of progeny virions^22^. However, the involvement of nuclear F-actin in the reorganization of marginalized chromatin caused by baculovirus remains undetermined. Here, we provided new insights into how baculovirus leverages the mechanism of heterochromatic organization to mediate chromatin marginalization. Lamin A/C-HP1a-H3K9me3 axis conferred a scaffold for the chromatin to localized at the nuclear periphery. Disrupting the association of Lamin A/C-HP1a-H3K9me3 axis affected the baculovirus induced chromatin marginalization and facilitated the nuclear egress of progeny virions. Moreover, we observed that the nuclear F-actin induced by baculovirus infection not only promotes the intranuclear transport of virions but also disturbs the nuclear actin pool, leading to the disassembly of chromatin. Our findings offer fundamental evidence for comprehending how DNA viruses manipulate host chromatin to modify the nuclear microenvironment.

## Results

### Characterization of the localization and interaction patterns between Lamin A/C and HP1a during viral infection

To better identify the baculovirus infection and track the single virion movement, we constructed a recombinant virus, named VP39-3mC, with *vp39* gene under its native promoter and fused to *3×mCherry* at its 3’ terminus, which is based on a *vp39* knockout bacmid preserved by our laboratory (Supplementary Fig. 1A). As a major capsid protein, VP39 mainly localizes at the VS and fused with 3×mCherry red fluorescent protein can be used to observe the motility of single virion in living cells^22^. VP39-3mC exhibits same replication and proliferation kinetics as the wild type virus (Supplementary Fig. 1B and 1C).

Next, we used VP39-3mC to explore the dynamic relationship between Lamin A/C and HP1a during viral infection. At 0 hours post infection (h p.i.), Lamin A/C localized at the nuclear membrane and showed no apparent location change during viral infection (Fig. 1A). As previous research reported, VP39-3mC infection also induced nuclear lamina disruption^23^ (Fig. 1D). HP1a was fused with a tagBFP fluorescent protein to indicate the localization of heterochromatin and Histone H2B was fused with a GFP fluorescent protein to indicate the localization of host chromatin. During VP39-3mC infection, HP1a gradually marginalized to the nuclear periphery and exhibited same location dynamics as the host chromatin (Fig. 1A). We observed a co-localization pattern between HP1a and Lamin A/C at 48 h p.i. (Fig. 1C), whereas this pattern was absent at 0 h p.i. (Fig. 1B). Additionally, they were not co-localized at the site where the nuclear lamina was disrupted (Fig. 1D), implying the integrity of nuclear lamina is critical for the heterochromatin marginalized to the nuclear periphery. The expression of Lamin A/C was down-regulated at the very late stage of infection, whereas HP1a was up-regulated at 24 and 48 h p.i. (Fig. 1E). These expression patterns indicated that HP1a may participate in the chromatin marginalization and Lamin A/C down-regulation may contribute to the disruption of nuclear lamina.

**Fig. 1.**
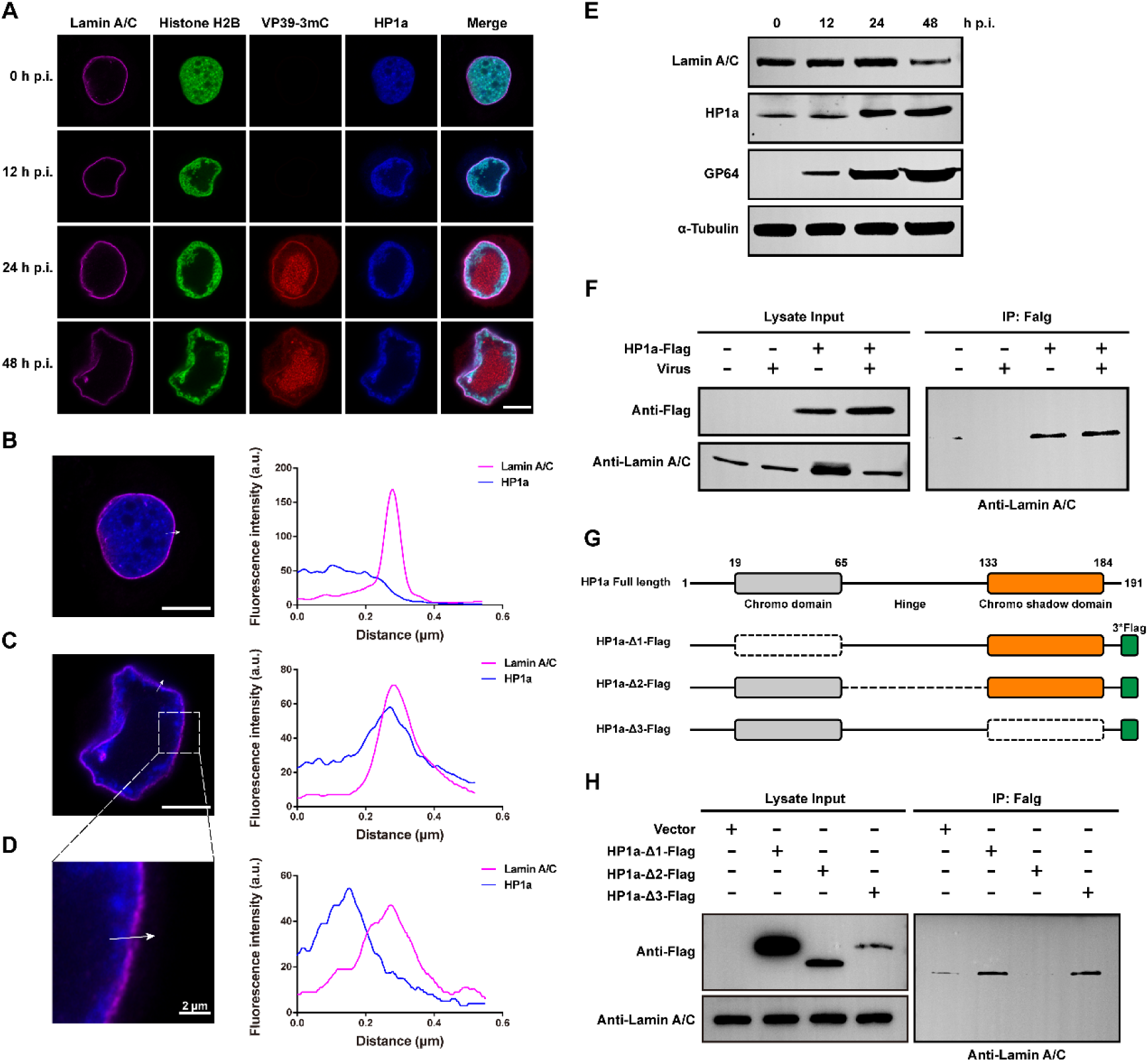
Colocalization and interaction between Lamin A/C and HP1a. **(A)** Immunofluorescence analysis by confocal microscopy. BmN cells were transfected with EGFP-Histone H2B (green) and tagBFP-HP1a (blue) plasmids and then infected with VP39-3mC virus (red). At indicated time point cells were incubated with anti-Dm0, a *drosophila* Lamin antibody, followed by treatment with secondary Alexa 647-conjugated antibody (purple). The presented cells are representative of a large number of cells. The scale bars represent 10 μm. **(B-D)** Colocalization analysis of Lamin A/C and HP1a at 0 h p.i. (**B**), 48 h p.i. (**C** and **D**). The scale bar represents 10 μm (**B** and **C**). **(E)** The expression pattern of Lamin A/C and HP1a during virus infection. **(F)** Co-IP assay analysis the interaction between Lamin A/C and HP1a. BmN cells were transfected with HP1a-Flag for 1 day and then infected or uninfected with WT virus for 48 h p.i. The cells were lysed and the proteins were immunoprecipitated with an anti-Flag monoclonal antibody. The precipitates were examined by western blot with a rabbit anti-Flag and anti-Lamin A/C polyclonal antibodies. **(G)** Schematic shows the domains of HP1a protein and the truncation mutants. **(G)** Co-IP assay analysis the domain of HP1a to mediate the interaction between Lamin A/C and HP1a. BmN cells were transfected with HP1a-Flag for 2 days and then lysed to extract total proteins. The proteins were immunoprecipitated with an anti-Flag monoclonal antibody. The precipitates were examined by western blot with a rabbit anti-Flag and anti-Lamin A/C polyclonal antibodies.

Then, we want to investigate whether in *Bombyx mori* Lamin A/C interacts with HP1a since it lacks *Lamin B* gene. The co-immunoprecipitation (Co-IP) assay revealed that transiently transfected HP1a-Flag interacted with endogenous Lamin A/C regardless of viral infection (Fig. 1F), suggesting the host heterochromatin localized at the nuclear periphery through Lamin A/C-HP1a interaction in the normal condition. *Bombyx mori* HP1a is a conserved protein with 191 aa (Supplementary Fig. 2A) and composed of a Chromo domain (N-terminal) and a Chromo shadow domain (C-terminal) connected by a Hinge region (Supplementary Fig. 2B). The Chromo domain and Chromo shadow domain mediate the recognition of H3K9me3. The Hinge region contained an intrinsically disordered region (Supplementary Fig. 2B), which is found to be important for the protein-protein interaction and phase separation^24, 25^. To identify the region responsible for the interaction between HP1a and Lamin A/C, we constructed three HP1a truncated mutants as shown in Fig. 1G. The Co-IP results indicated that truncating the Hinge region of HP1a eliminates the interaction between Lamin A/C (Fig. 1H).

Taken together, our findings suggested that in *Bombyx mori* HP1a interacts with Lamin A/C by its Hinge region and this interaction may participate in the baculovirus infection induced chromatin marginalization.

### Lamin A/C-HP1a interaction mediates the baculovirus infection induced chromatin marginalization

To elucidate the function of Lamin A/C-HP1a interaction, we knocked down (KD) the expression of Lamin A/C and HP1a by using RNAi (Supplementary Fig. 3A and 3B). Knocking down Lamin A/C (Lamin A/C-KD) alone or simultaneously knocking down both Lamin A/C and HP1a (Double-KD) showed no significant effects on viral replication and proliferation (Supplementary Fig. 3C and 3D). In control-KD cells, nuclear lamina remained intact with the host chromatin tightly localized at the nuclear periphery (Fig. 2A). In Lamin A/C-KD cells, however, nuclear lamina was disrupted leaving only an irregular fraction within the nucleus (Fig. 2A), which is consistent with our previous report^26^. Silencing of Lamin A/C resulted in the host chromatin primarily localizing at nuclear lamina fragment, altering the marginalization of chromatin caused by viral infection (Fig. 2A). Furthermore, silencing of both Lamin A/C and HP1a not only disrupted the nuclear lamina but also resulted in the dispersed distribution of the host chromatin in the nucleus (Fig. 2A). To analysis the disruption of chromatin marginalization, we compared the proportion of nuclear area occupied by chromatin during viral infection. As shown in Supplementary Fig. 3E, baculovirus infection induced a progressive decrease in the ratio of chromatin area in the nucleus, indicating the chromatin marginalization. siRNA-mediated KD of Lamin A/C partially inhibited viral infection induced chromatin marginalization (Fig. 2B). Similarly, Double-KD also increased the ratio of chromatin area at 24 and 48 h p.i. (Fig. 2B). This defect was rescued by co-overexpressing Lamin A/C and full-length HP1a, but not by overexpressing the Hinge-deleted mutant of HP1a (Fig. 2B). Thus, Lamin A/C-HP1a interaction is involved in the baculovirus infection induced chromatin marginalization.

**Fig. 2.**
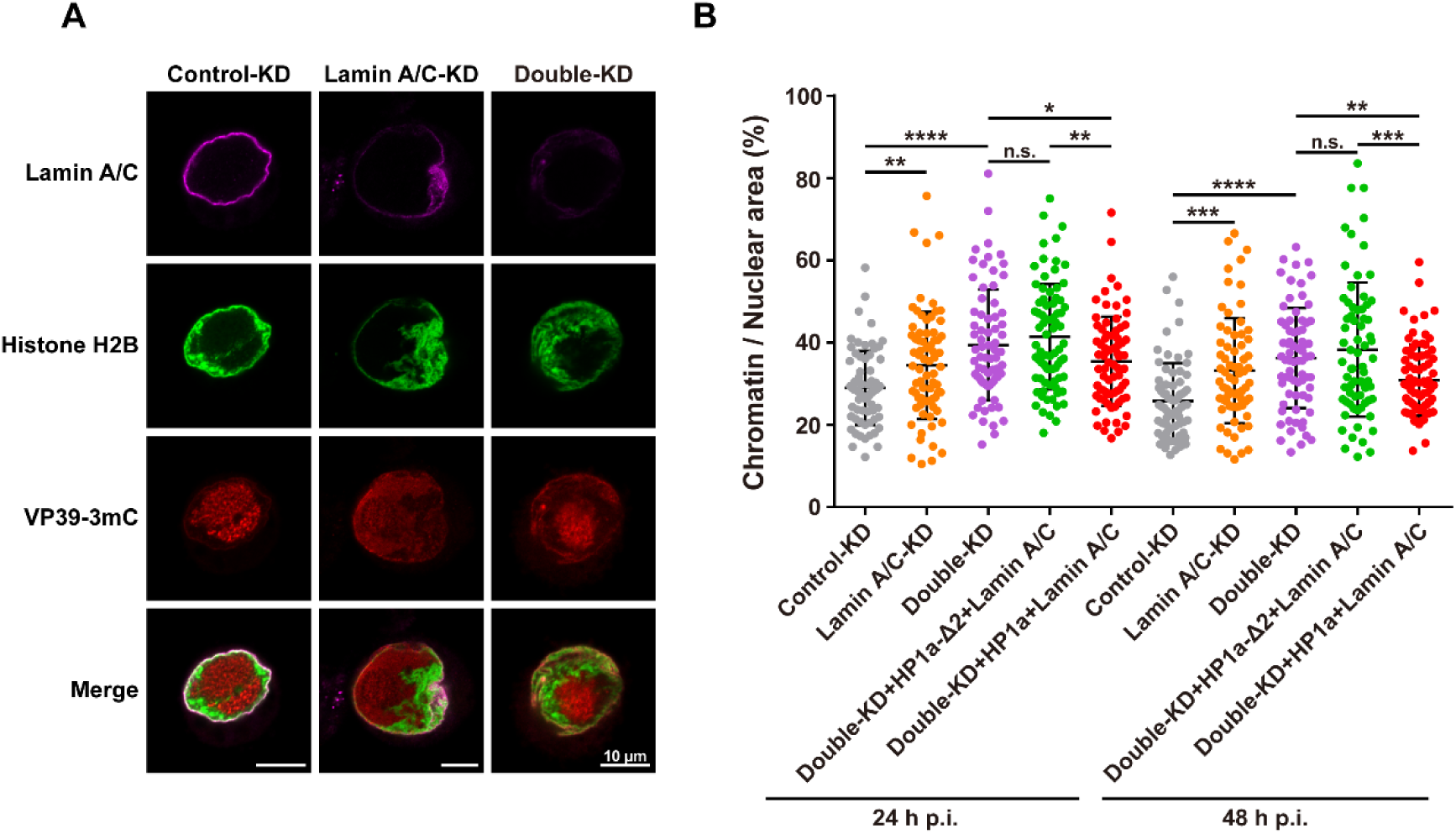
The functions of Lamin A/C and HP1a in chromatin marginalization. **(A)** Silencing of Lamin A/C and HP1a impaired the chromatin marginalized to the nuclear periphery. BmN cells were transfected with EGFP-Histone H2B (green) plasmid for 1day and then transfected with siRNAs for 1 day. The cells were infected with VP39-3mC virus (red). At 48 h p.i., the cells were incubated with anti-Dm0 followed by treatment with secondary Alexa 647-conjugated antibody (purple). The presented cells are representative of a large number of cells. The scale bars represent 10 μm. **(B)** Quantitation analysis of the ratio of chromatin area in the nucleus. BmN cells were transfected with siRNAs for 1 day and then transfected with EGFP-Histone H2B, Lamin A/C-HA and HP1a-Δ2-Flag or HP1a-Flag for 1 day. The cells were infected with VP39-3mC virus at 24 and 48 h p.i. (n > 60). Error bars indicate mean ± SD. Student’s *t*-test was performed, *, P < 0.05; **, P < 0.01; ***, P < 0.001; ****, P < 0.0001; n.s., no significance.

### Lamin A/C-HP1a-H3K9me3 axis mediates the chromatin marginalization induced by baculovirus

As a well-known heterochromatic protein, HP1a can recognize and bind Histone H3 tri-methyl K9 (H3K9me3) modification. Therefore, we next aimed to investigate whether H3K9me3 participates in the viral infection induced chromatin marginalization. During VP39-3mC infection, the location of H3K4me3 (a euchromatic marker) was marginalized with the host chromatin, while H3K9me3 exhibited a distinct localization pattern (Fig. 3A and 3B). Specifically, H3K9me3 was mainly located at the nuclear periphery at 0 h p.i., but its location altered to colocalize with the Histone H2B at 12 h p.i. and gradually marginalizing with the host chromatin at 24 and 48 h p.i. (Fig. 3B and 3C). This translocation indicated that H3K9me3 might contribute to the host chromatin marginalization during viral infection. Additionally, we tested the impact of viral infection on the binding of HP1a to H3K9me3. Strikingly, our Co-IP results demonstrated that HP1a was able to bind H3K9me3 after viral infection, whereas lacked the ability to bind H3K9me3 in the absence of viral infection (Fig. 3D).

**Fig. 3.**
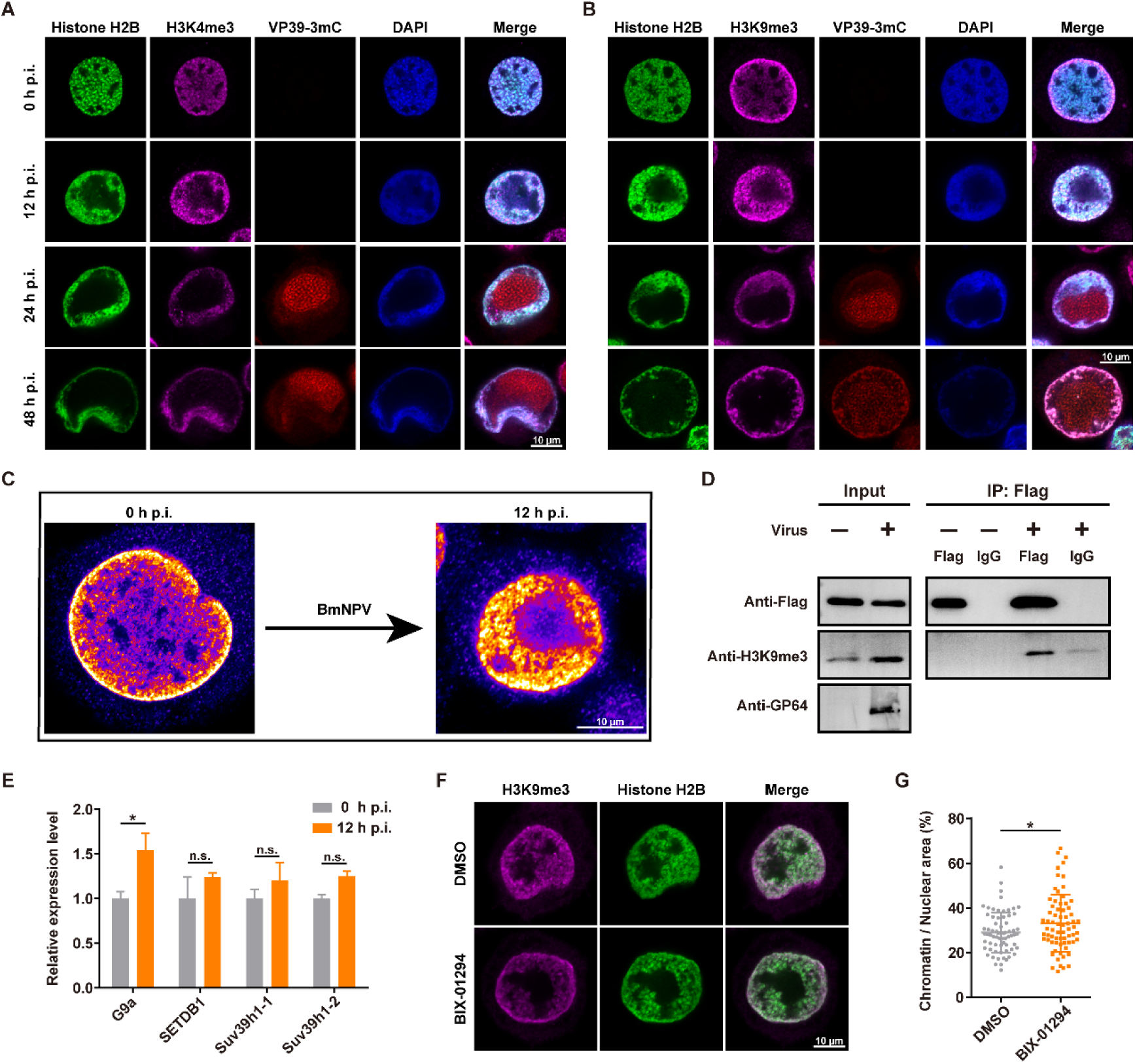
H3K9me3 participates in the Lamin A/C and HP1a mediated chromatin marginalization. **(A and B)** The location of H3K4me3 (A) and H3K9me3 (B) during baculovirus infection. The BmN cells were transfected with EGFP-Histone H2B for 1 day and then infected with VP39-3mC virus. At indicated time points, the cells were incubated with anti-H3K4me3 or anti-H3K9me3 antibody followed by treatment with secondary Alexa 647-conjugated antibody (purple). DAPI was used to stain the DNA (blue). The presented cells are representative of a large number of cells. The scale bars represent 10 μm. **(C)** The location transition of H3K9m3 at the early stage of infection. The scale bars represent 10 μm. **(D)** Co-IP analysis the interaction between HP1a and H3K9me3. BmN cells were transfected with HP1a-Flag plasmid for 1 day and then infected or uninfected with WT virus for 2 days. **(E)** Changes in HMTs transcription (measured by qRT-PCR) at 0 and 12 h p.i. (n = 3) **(F)** The effect of BIX-01294 treatment on H3K9me3 location transition during viral infection. The scale bars represent 10 μm. **(G)** The effect of BIX-01294 treatment on the chromatin marginalization at 24 h p.i. (n > 60). Error bars indicate mean ± SD. Student’s *t*-test was performed, *, P < 0.05; n.s., no significance.

These findings guide us to further investigate the factors responsible for the translocation of H3K9me3 at 12 h p.i. Methylation of H3K9 is mediated by histone methyltransferases (HMTs)^27^, therefore, we examined the expression of HMTs in response to viral infection. The qRT-PCR results suggested that only the expression of G9a was increased at 12 h p.i. (Fig. 3E), indicating this HMT is a potential candidate for mediating the H3K9me3 translocation. To address this possibility, a specific inhibitor, BIX-01294, was used to study the function of G9a. The efficiency of BIX-01294 in BmN cells and its effects on viral replication and proliferation have been validated (Supplementary Fig. 4). BIX-01294 treatment resulted in significant defects in H3K9me3 translocation (Fig. 3F) and chromatin marginalization (Fig. 3G) during viral infection. Thus, our findings revealed that G9a mediated H3K9me3 translocation and the increased binding of HP1a to H3K9me3 promote the marginalization of chromatin.

### Marginalized chromatin impedes the nuclear export of progeny virions

Generally, it is widely believed that viruses remodel the nuclear microenvironment to enhance their replication and proliferation. Recently, however, several researches suggested that HSV-1 infection induced marginalized chromatin physically restricted the diffusion of its capsids in the nucleus^5, 28^. While HSV-1 capsids move through diffusion^29^, baculovirus capsids transport within the nucleus by utilizing nuclear F-actin^22^. Given the distinct nuclear motion types and different virion morphologies between these two viruses^6, 30^, we want to reveal whether the nuclear export of baculovirus is also hindered by the marginalized chromatin. VP39-3mC was used to monitor single virion motion by using live cell imaging. In live BmN cells, we observed a single virion transport was impeded by the marginalized chromatin during nuclear export (Fig. 4A and Video 1). This collision was also monitored in other cells (Video 2 and 3), indicating it is not a random event occurring in the nucleus and chromatin marginalization indeed physically inhibits the nuclear export of baculovirus. To quantitatively analyze this obstruction caused by marginalized chromatin, the efficiency of virus budding was studied by measuring viral genome copies in the supernatant. As shown in Fig. 4B, the highest budding efficiency between 36-48 h p.i. and gradually decreased thereafter, which coincided with the time period of chromatin marginalization. Based on the above results, Double-KD was introduced to disrupt chromatin marginalization. In Double-KD cells, the levels of viral genome copies in the cytoplasm were significantly higher than in the control (Fig. 4C), suggesting marginalized chromatin physically impedes the nuclear export of progeny virions. This change in nuclear export was not due to the reduction in the motion ability of progeny virions at the very late stage of infection (Video 4). Taken together, we proposed that baculovirus infection induces chromatin marginalization to create space for the formation of VS and assembly of nucleocapsids, which in turn impeding the nuclear export of progeny virions. Thus, a substantial abundance of virions was retained in the nucleus to generate ODVs and subsequently embedded into OBs.

**Fig. 4.**
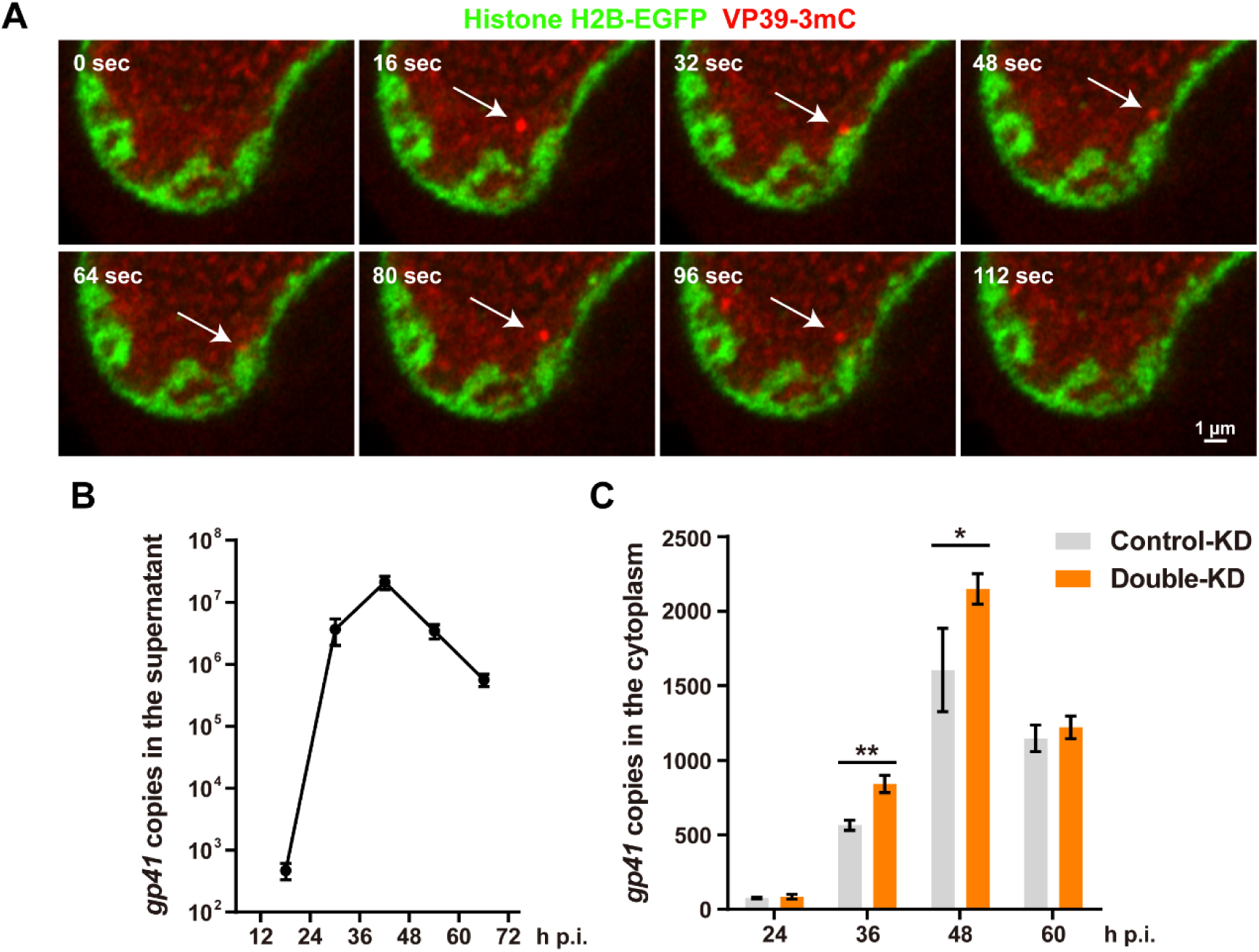
Live cell imaging analysis the role of marginalized chromatin in nuclear export of progeny virions. **(A)** Time series of a single virion movement at 46 h 35m p.i. The BmN cells were transiently transfected with EGFP-Histone H2B plasmid and then infected with VP39-3mC virus. White arrows indicate the single virion. The scale bar represents 1 μm. **(B)** The viral genome copies in the supernatant within a 12-hour interval (n = 3). **(C)** The viral genome copies in the cytoplasm after virus infection (n = 3). Error bars indicate mean ± SD. Student’s *t*-test was performed, *, P < 0.05; **, P < 0.01.

### Nuclear F-actin mediates alteration in chromatin organization through disrupting the dynamics of nuclear actin pool

Recently, we reported that the host chromatin genome-widely gained accessibility is accompanied by the disassembly of multi-nucleosome in response to baculovirus infection^4^. The mechanism underlying changes in chromatin organization is still unclear. After mitosis, the dynamics of nuclear actin pool controls chromatin organization and a cell-cycle-specifically formed nuclear F-actin reorganizes the chromatin architecture^20^. This prompted us to unveil the role of baculovirus infection induced nuclear F-actin in altering the chromatin organization. To label the nuclear F-actin, we used our laboratory previously constructed vector, Lifeact-EGFP^26^, to transiently express Lifeact fused with a EGFP fluorescent protein and a nuclear localization signal (NLS). At 24 h p.i., a large number of actin filaments were observed within the nucleus and these actin filaments disappeared at 48 h p.i. (Fig. 5A). Subsequently, fluorescence recovery after photobleaching (FRAP) was employed to identify the dynamics of nuclear actin pool. We transiently transfected a plasmid to express wild-type actin fused with EGFP and NLS. The time of recovery for wild-type actin at 0 h p.i. was greater than at 24 h p.i. (Fig. 5B), suggesting viral infection disrupt the dynamics of nuclear actin pool. Furthermore, R62D, a polymerization-deficient actin mutant, was transfected to dilute the amount of actin monomers within the nucleus and rescued the recovery time of nuclear actin (Fig. 5B).

**Fig. 5.**
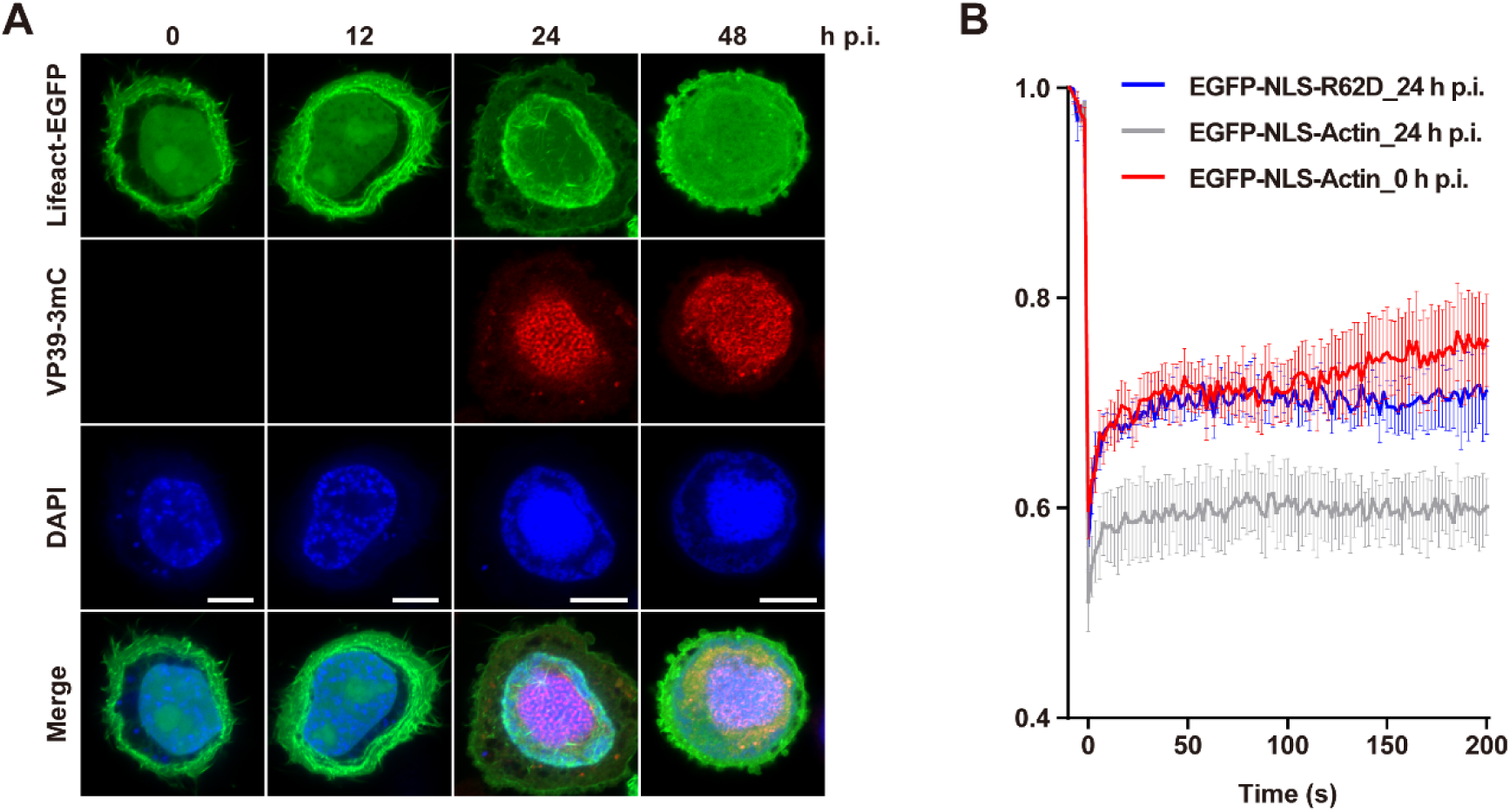
Baculovirus infection disrupts the dynamic of nuclear actin pool. **(A)** The formation of nuclear F-actin during baculovirus infection. The BmN cells were transfected with Lifeact-EGFP-NLS plasmid and then infected with VP39-3mC virus. DAPI was used to stain the DNA. The presented cells are representative of a large number of cells. The scale bars represent 10 μm. **(B)** FRAP assay analysis of the recovery of the G-actin in the nucleus during viral infection.

Quantification analysis of chromatin density by Histone H2B-EGFP fluorescence intensities revealed a significant reduction in chromatin compaction at 24 h p.i. (Fig. 6A and 6B). To determine the importance of nuclear actin polymerization in chromatin decompaction during viral infection, the cells were treated with drug that prevent actin polymerization (Cytochalasin D, CD) at 12 h p.i. The H2B intensities were rescued by CD treatment (Fig. 6C and 6D). Notwithstanding, we could not conclude that the different observation in chromatin compaction is mediated by nuclear F-actin, because CD could induce actin depolymerization both in the cytoplasm and the nucleus. Hence, mCherry-NLS-R62D was transiently expressed to rescue the dynamics of nuclear actin pool and inhibit the formation of nuclear actin filaments. As expected, mCherry-NLS-R62D also could prevent the chromatin decompaction during viral infection (Fig. 6E and 6F).

**Fig. 6.**
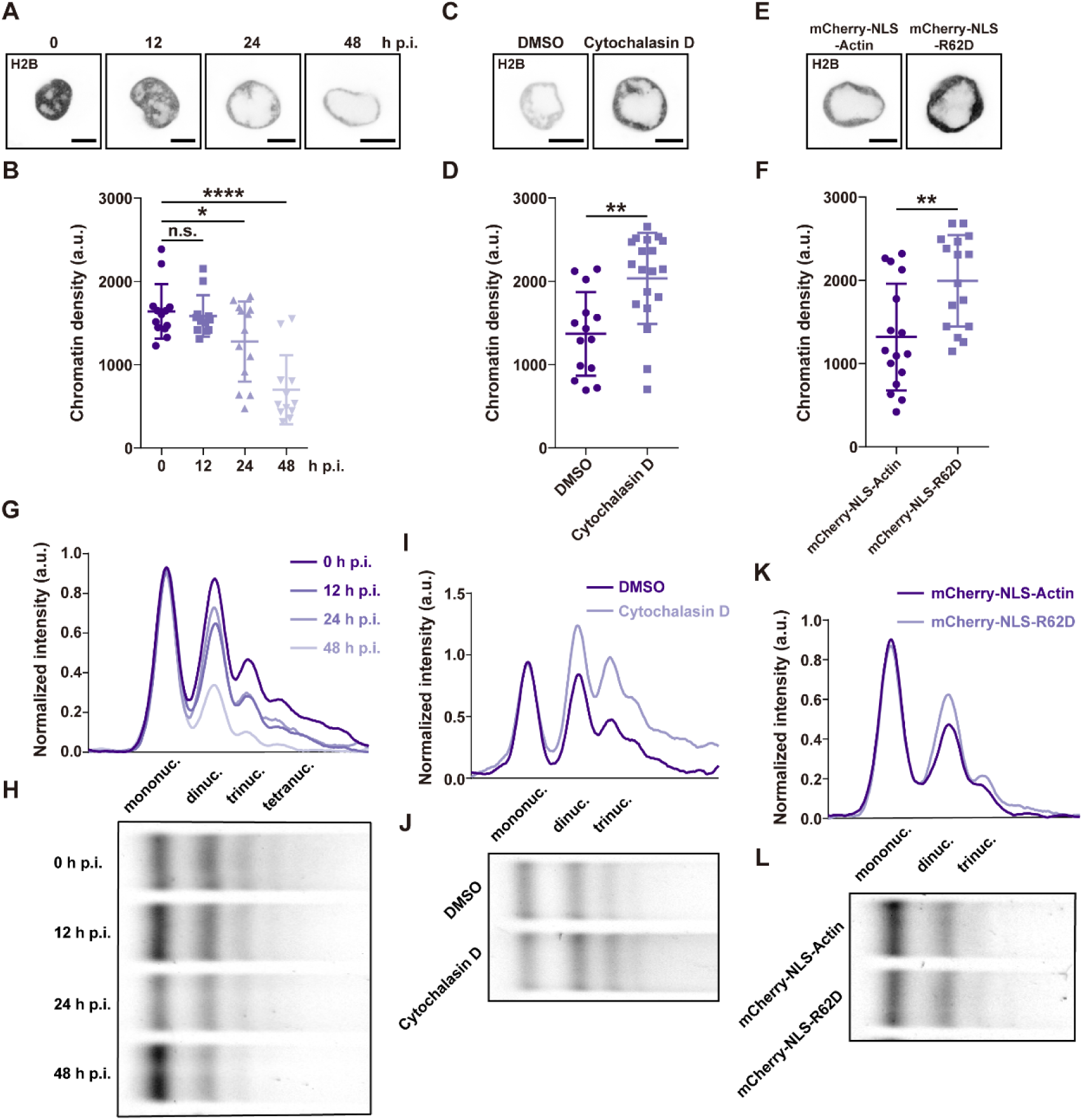
Nuclear F-actin involves in the viral infection induced chromatin architectural changes. **(A)** The EGFP-Histone H2B fluorescence densities (grey) during viral infection. **(B)** Chromatin densities of BmN cells at indicated times of infection (n = 12 or 13). **(C)** The EGFP-Histone H2B fluorescence densities with the treatment of DMSO or CD at 24 h p.i. **(D)** Chromatin densities of BmN cells with the treatment of DMSO or CD at 24 h p.i. (n = 14 (DMSO) or 19 (CD)). **(E)** The EGFP-Histone H2B fluorescence densities with the transfection of mCherry-NLS-Actin or mCherry-NLS-R62D at 24 h p.i. **(F)** Chromatin densities of BmN cells with the transfection of mCherry-NLS-Actin or mCherry-NLS-R62D at 24 h p.i. (n = 16). **(G** and **H)** Analysis of chromatin compaction by MNase accessibility assay at indicated times of infection. **(I** and **J)** Analysis of chromatin compaction by MNase accessibility assay with the treatment of DMSO or CD at 24 h p.i. **(K** and **L)** Analysis of chromatin compaction by MNase accessibility assay with the transfection of mCherry-NLS-Actin or mCherry-NLS-R62D at 24 h p.i. The scale bar represents 10 μm. Error bars indicate mean ± SD. Student’s *t*-test was performed, *, P < 0.05; **, P < 0.01; ****, P < 0.0001; n.s., no significance.

To more directly confirm the role of nuclear actin in chromatin organization, MNase digestion was performed to measure the nucleosome assembly^31^. Consistent with us previous findings, baculovirus infection resulted in multi-nucleosome disassembly (Fig. 6G and 6H). Moreover, CD treatment or expressing mCherry-NLS-R62D appeared more resistance to MNase digestion compared with DMSO or mCherry-NLS-Actin (Fig. 6I-L), indicating the formation of nuclear F-actin facilitates the chromatin decompaction during viral infection.

Taken together, these findings provided a connection between the dynamics of nuclear actin and chromatin organization during viral infection. Baculovirus infection induces the polymerization of nuclear actin, thereby disrupting the dynamics of nuclear actin pool and resulting in chromatin decompaction as well as multi-nucleosome disassembly.

## Discussion

With the development of genomic sequencing and technologies, there is increasing interest in understanding how viruses manipulate the host chromatin. In this work, we elucidated the mechanistic basis for chromatin marginalization and architectural alteration during baculovirus infection. Lamin A/C-HP1a-H3K9me3 axis mediates the chromatin marginalized to the nuclear periphery and marginalized chromatin impedes the nuclear export of progeny virions. Moreover, baculovirus infection induced nuclear F-actin is a contributing factor to chromatin decompaction and nucleosome depletion.

The replication of viral genome and the formation of VRC are considered as the main factors inducing the host chromatin marginalization^32, 33^. However, why host chromatin does not exhibit polarity localization like in Human Cytomegalovirus (HCMV) infection and instead distributes at the nuclear periphery has remained largely unexplained. In fact, the phenomenon of chromatin marginalization also exists in mammals. H3K9me3 is associated with the nuclear lamina through interacting with Lamin B-LBR-HP1a^11, 34^. We found that this conserved mechanism is also exploited by baculovirus to mediate the whole genome marginalized to the nuclear lamina. The crucial step in this process is the G9a-mediated repositioning of H3K9me3, which leads to H3K9me3 localized in the nucleoplasm providing more targets for HP1a recognition.

Recently, a study has attempted to explore the significance of Lamin A/C and LBR on HSV-1 infection^3^. They found that Lamin A/C and LBR knockout impaired HSV-1 genes expression, as well as the formation of VRC and the marginalization of chromatin. Owing to the compromised HSV-1 genes expression and VRC formation, it is challenging to straightforwardly confirm how Lamin A/C and LBR prevent the chromatin marginalization. Here, knockdown of Lamin A/C and HP1a in BmN cells has no influence on the baculovirus replication and proliferation. Moreover, silkworm provides a relatively simple genetic background to explore the function of Lamin A/C mediated chromatin marginalization during baculovirus infection.

Our findings in live cell imaging strongly support the hypothesis that nuclear egress of progeny virions is hindered by marginalized chromatin. This phenomenon is conserved in both HSV-1 and baculovirus, even though they have different virion structure. At the very late stage of infection, BV production is shifted to the retention of nucleocapsids in the nucleus and nucleocapsids are incorporated into occlusion bodies^35^. Only ∼2.3% progeny virions are budded out of infected cells, while the most of them are retained in the nucleus^36, 37^. The transition from BV to ODV production is thought to be the result of ODV associated proteins accumulate in the nucleus^38–43^. Once the nucleocapsids are enveloped by intranuclear vesicles, they will no longer transit to the cytoplasm. Our results shed novel insights into the shift of BV to ODV production, which could be the result of physical changes in the nuclear microenvironment at the very late stage of infection, providing a new direction for understanding the origin of the biphasic life cycle of baculovirus.

Furthermore, we also reveal the role of baculovirus infection induced nuclear F-actin in the changes in chromatin architecture. The formation of nuclear actin filaments disturbs the balance of actin pool, thereby causing the increased accessibility of chromatin and disassembly of multi-nucleosome. These observations are similar with the findings during mitosis exit. Despite we have linked nuclear F-actin to changes in chromatin architecture, the molecular mechanism involved in this process remain elusive. Nuclear actin plays a vital role as a key component in several chromatin remodeling complexes^15, 44, 45^. Recent research has found that the loss of nuclear actin leads to chromatin remodeling, which is mediated through the dysregulation of BAF chromatin remodeling complex^44^. These findings will guide our further exploration into whether the formation of nuclear actin filaments, which disrupts the dynamics of nuclear actin pool, also affect the functionality of chromatin remodeling complexes to reorganize the chromatin architecture.

Overall, we characterized the molecular mechanism underlying the chromatin marginalization and decompaction during baculovirus infection. Our findings revealed that the marginalized chromatin functions as a physical barrier to impede the nuclear export of progeny virions. These results are essential in comprehending the strategies employed by DNA viruses to manipulate the host nuclear microenvironment.

## Materials and Methods

### Cell culture, virus strain, antibodies and plasmids

BmN cells were maintained at 27 ℃ as a monolayer in the Sf-900 II SFM medium (Invitrogen) supplemented with 3% fetal bovine serum (FBS, Invitrogen). The T3 strain of BmNPV was termed as a wild-type (WT) virus. The recombinant virus used in this work VP39-3mC was constructed via RecE/RecT (ET) homologous recombinantion and T7-mediated transposition, which contained the *vp39-3×mCherry* driven by *vp39* native promoter.

The rabbit anti-Lamin A/C polyclonal antibody was a kind gift of Prof. Qingyou Xia (Southwest University, Chongqing, China). Commercially available antibodies against HP1a (Abcam, ab109028), H3K4me3 (Abcam, ab8580), H3K9me3 (Abcam, ab8898), Dm0 (DSHB, ADL67.10), α-Tubulin (Beyotime, AF0001), Flag tag (HUABIO, 0912-1) and GP64 (Invitrogen, 14-6995-85) were used.

The transient expression plasmids including EGFP-Histone H2B, tagBFP-HP1a, Lifeact-EGFP-NLS, EGFP-NLS-Actin, EGFP-NLS-R62D, mCherry-NLS-Actin, mCherry-NLS-R62D, HP1a-Flag, HP1a-Δ1-Flag, HP1a-Δ2-Flag, HP1a-Δ3-Flag and Lamin A/C-HA were generated from the pIZ/V5-His plasmid and preserved by our laboratory.

### Immunofluorescence (IF)

BmN cells on the 35 mm coverslips were transfected with the expression plasmids using the Lipo8000 Transfection Reagent (Beyotime, C0533) according to the manufacturer’s protocol and then infected with VP39-3mC virus after 2 days post-transfection at an MOI of 10. Cells were fixed with 4% paraformaldehyde for 15 min, permeabilized with 0.1% TritonX-100 for 20 min, blocked with 5% BSA/PBS for 2 h, and incubated with primary antibodies anti-Dm0 (dilution 1:20), anti-H3K4me3 (dilution 1:200) and anti-H3K9me3 (dilution 1:500) overnight at 4 ℃. Afterwards, cells were incubated with secondary Alexa 647-conjugated donkey anti-rabbit or anti-mouse antibody (Beyotime) according to the experiments. Then, cells were stained with DAPI (Beyotime) and examined through ZEISS LSM 880 confocal scanning laser microscopy.

### Drug treatments

For the inhibitor treatments, BIX-01294 (MCE, HY-10587, 20 and 40 µM) was added into the medium at 0 h p.i. and Cytochalasin D (Abcam, ab143484, 1µg/ml) was added at 12 h p.i.

### Immunoblotting

For immunoblotting assay, 30 µg total proteins at indicated time points and treatments were mixed with 2×Laemmli buffer, separated through 12% or 15% SDS-PAGE, and examined through western blotting using the rabbit anti-Lamin A/C polyclonal antibody (dilution 1:5000), mouse anti-GP64 monoclonal antibody (dilution 1:5000), rabbit anti-α-Tubulin polyclonal antibody (dilution 1:1000), rabbit anti-Flag polyclonal antibody (dilution 1:5000), rabbit anti-H3K9me3 polyclonal antibody (dilution 1:2000), rabbit anti-HP1a monoclonal antibody (dilution 1:1000).

### Co-immunoprecipitation (Co-IP)

For Co-IP assay, BmN cells were transfected with HP1a-Flag/HP1a-Δ1-Flag/HP1a-Δ2-Flag/HP1a-Δ3-Flag expression plasmids. After 1 day post-transfection, cells were infected with WT virus at an MOI of 10. Infected cells were collected at 2 days post infection and washed three time with PBS supplemented with protease inhibitor cocktail (Bimake, B14001). The cell pellets were resuspended in lysis buffer (Beyotime, P0013) supplemented with protease inhibitor cocktail. Then, cell lysates were incubated with mouse anti-Flag immunomagnetic beads (Bimake, B26101) overnight at 4 ℃. Afterwards, beads were harvested with a magnetic separator and washed with PBS supplemented with protease inhibitor cocktail five times. The samples were subjected to SDS-PAGE and examined by immunoblotting.

### RNA interference

The control siRNA and siRNAs targeted the host *Lamin A/C* and *HP1a* genes were synthesized from Sangon Biotech. The siRNAs were transfected using LipoRNAi Transfection Reagent (Beyotime, C0535) according to the manufacturer’s protocol.

The sequences of siRNA were shown as follows: Control-siRNA (Sense: UUC UCC GAA CGU GUC ACG UTT; Antisense: ACG UGA CAC GUU CGG AG AATT); *Lamin A/C*-siRNA1 (Sense: GCA AGA GAA AGA UGC AUU ATT; Antisense: UAA UGC AUC UUU CUC UUG CTT); *Lamin A/C*-siRNA2 (Sense: GCG AAC AAC AGG AGG CAA ATT; Antisense: UUU GCC UCC UGU UGU UCG CTT); *Lamin A/C*-siRNA3 (Sense: GGA AGU CGC AAU UUC UGA ATT; Antisense: UUC AGA AAU UGC GAC UUC CTT); *HP1a*-siRNA1 (Sense: AAG AAU AUG UCG UGG AGA ATT; Antisense: UUC UCC ACG ACA UAU UCU UTT); *HP1a*-siRNA2 (Sense: GGG AAA GAA AGG AAG AAA ATT; Antisense: UUU UCU UCC UUU CUU UCC CTT); *HP1a*-siRNA3 (Sense: AAA CUU AGA CUG UGA GGA ATT; Antisense: UUC CUC ACA GUC UAA GUU UTT);

### Quantitative real-time PCR (qRT-PCR) analysis

BmN cells infected with WT virus at indicated time were harvested and total RNA was extracted by using TRIzol reagent (Invitrogen). 2 μg total RNA was prepared to synthesis the cDNA by TransScript One-Step gDNA Removal and cDNA Synthesis SuperMix (TransGen). qRT-PCR was employed using Hieff qPCR SYBR Green Master Mix (Yeasen) and the primers of qRT-PCR were used as follows: *G9a*-F: 5’-CAC GGA GGT CGA GAC GAT TA-3’, *G9a*-R: 5’-TGC TGG CTC TGC TTT GAC TT-3’; *SETDB1*-F: 5’-TTG GAC GGA CTC AAC GAA GG-3’, *SETDB1*-R: 5’-TTC CTG CTT CAC ACC ACC AG-3’; *Suv39h1-1*-F: 5’-AAG GCA GGC GAC GAT TAA CA-3’, *Suv39h1-1*-R: 5’-GAA TCC GAC AGG CGC TCT AA-3’; *Suv39h1-2*-F: 5’-TGC TCT CTC TCC CGA AGT CA-3’, *Suv39h1-2*-R: 5’-TCC GAC AGG CGC TCT AAT TT-3’.

### Live cell imaging

For live cell imaging to visualize single virion motion, BmN cells on 35 mm coverslips were transfected with EGFP-Histone H2B plasmid. After one day post transfection, cells were infected with VP39-3mC at an MOI of 20. The images were captured at 1 s intervals between 46 to 48 h p.i. using the Airyscan imaging system of ZEISS LSM 880 confocal scanning laser microscopy.

### Analysis of viral growth curve

BmN cells were infected with WT or VP39-3mC at an MOI of 10. At the indicated time points, supernatants containing the BV were harvested, and cell pellets were removed by centrifugation at 3000 rpm for 5 min. The titers of BV were determined by TCID_50_ end-point dilution assay in 96 wells plate.

### Quantification of viral genome copies

For analysis of the viral genome replication, the total DNA were extracted using TaKaRa MiniBEST Universal Genomic DNA Extraction Kit. qPCR analysis of viral genomic DNA was performed with *gp41* primers (F: 5’-GCA CAT CAA CAT GAT CAA CG-3’, R: 5’-TAA ACT CAT GAT TCG CGC TC-3’).

For analysis of the viral genome copies in the supernatants, the viral genomic DNA in the supernatants were extracted using TaKaRa MiniBEST Viral RNA/DNA Extraction Kit. qPCR analysis of viral genomic DNA was performed with *gp41* primers. For analysis of the viral genome copies in the cytoplasm, the components of cytoplasm were extracted using Buffer A (10 mM HEPES, 2 mM MgCl_2_, 1 × protease inhibitor cocktail and 0.6 % NP-40), the components of nucleus were removed by centrifugation at 1000 g for 10 min. Then, the viral genomic DNA was extracted using TaKaRa MiniBEST Viral RNA/DNA Extraction Kit. qPCR analysis of viral genomic DNA was performed with *gp41* primers.

### Micrococcal Nuclease (MNase) digestion assay

Five million cells were harvested and washed once with PBS. After centrifugation at 3000 rpm for 5 min, the cell pellet was resuspended in 1 mL Buffer A mentioned above and homogenized. Nuclei were collected by centrifugation at 1000 g for 10 min and washed once with Buffer A. Then, nuclei were resuspended with 500 μL MNase Buffer (50 mM Tris-HCl, pH 7.9, and 5 mM CaCl_2_). MNase digestion was performed by addition of 1 μL MNase (NEB, M0247S) at 37 ℃ for 5 min. Reaction was terminated by adding 1/10 (v/v) EGTA (0.5 M), 1/10 (v/v) EDTA (0.5 M), 1/100 (v/v) SDS (10 %), 5 μL Proteinase K and 2.5 μL RNase A at 56 ℃ for 15 min. DNA was purified using a PCR purification kit and 500 ng of DNA was analysis on a 1.5 % agarose gel.

### Fluorescence recovery after photobleaching (FRAP) assay

For FRAP experiments, BmN cells were plated on 35 mm coverslips and transfected with EGFP-NLS-Actin or EGFP-NLS-R62D. After one day post transfection, cells were infected with WT virus. At indicated time points, FRAP was performed using the FRAP module of ZEISS LSM 880 confocal scanning laser microscopy.

### Statistical analysis

Student’s t-tests were conducted using the GraphPad Prism6. p < 0.05 indicated statistical significance.

## Acknowledgments

We sincerely extend our deep thanks to Prof. Qingyou Xia (Southwest University, China) for the rabbit polyclonal anti-Lamin A/C antibody. We acknowledge Yunqing Li (Zhejiang University, China) for technical assistance on live cell imaging. We are also grateful to the Dr. Taishu Wang (Dalian Medical University) for the pEGFP-C1-Lifeact plasmid.

## Funding Information

This work was supported by the National Natural Science Foundation of China (32172793/31972619), the Natural Science Foundation of Zhejiang Province (Z20C170008) and Postdoctoral Research Foundation of China (2021M702871). The funders had no role in study design, data collection and analysis, decision to publish, or preparation of the manuscript.

## Author Contributions

**Conceptualization:** Xiangshuo Kong, Xiaofeng Wu.

**Investigation:** Xiangshuo Kong, Guanping Chen, Conghui Li.

**Methodology:** Xiangshuo Kong, Conghui Li, Xiaofeng Wu.

**Supervision:** Xiaofeng Wu.

**Visualization:** Xiangshuo Kong, Guanping Chen.

**Writing – original draft:** Xiangshuo Kong.

**Writing – review & editing:** Xiaofeng Wu.

## Disclosure Declaration

The authors declare no competing financial interests.

## Abbreviations

The following abbreviations are used in this manuscript:

BmNPV: Bombyx mori nucleopolyhedrovirus NL nuclear lamina
HP1a: heterochromatin protein 1 alpha
ODV: occlusion-derived virion
OB: occlusion body
BV: budded virion
VRC: viral replication compartment
HSV-1: herpes simplex virus 1
HCMV: human cytomegalovirus
VS: virogenic stroma
LBR: Lamin B receptor
HMTs: histone methyltransferases
CD: Cytochalasin D
FRAP: fluorescence recovery after photobleaching MOI multiplicity of infection
h p.i.: hours post-infection

## Notes

### Competing Interest Statement

The authors have declared no competing interest.

## References

1. Knipe, D.M., Prichard, A., Sharma, S. & Pogliano, J. Replication Compartments of Eukaryotic and Bacterial DNA Viruses: Common Themes Between Different Domains of Host Cells. Annu Rev Virol 9, 307–327 (2022).

2. Paulus, C., Nitzsche, A. & Nevels, M. Chromatinisation of herpesvirus genomes. Reviews in medical virology 20, 34–50 (2010).

3. Takeshima, K., Maruzuru, Y., Koyanagi, N., Kato, A. & Kawaguchi, Y. Redundant and Specific Roles of A-Type Lamins and Lamin B Receptor in Herpes Simplex Virus 1 Infection. J Virol 96, e0142922 (2022).

4. Kong, X. et al. Dynamic chromatin accessibility profiling reveals changes in host genome organization in response to baculovirus infection. PLoS Pathog 16, e1008633 (2020).

5. Aho, V. et al. Infection-induced chromatin modifications facilitate translocation of herpes simplex virus capsids to the inner nuclear membrane. PLoS Pathog 17, e1010132 (2021).

6. Rohrmann, G.F. Baculovirus Molecular Biology, 2019.

7. Nagamine, T., Kawasaki, Y., Abe, A. & Matsumoto, S. Nuclear marginalization of host cell chromatin associated with expansion of two discrete virus-induced subnuclear compartments during baculovirus infection. J Virol 82, 6409–6418 (2008).

8. Blissard, G.W. & Theilmann, D.A. Baculovirus Entry and Egress from Insect Cells. Annu Rev Virol 5, 113–139 (2018).

9. Wong, X., Melendez-Perez, A.J. & Reddy, K.L. The Nuclear Lamina. Cold Spring Harbor perspectives in biology 14 (2022).

10. Pindyurin, A.V. et al. The large fraction of heterochromatin in Drosophila neurons is bound by both B-type lamin and HP1a. Epigenetics & chromatin 11, 65 (2018).

11. van Steensel, B. & Belmont, A.S. Lamina-Associated Domains: Links with Chromosome Architecture, Heterochromatin, and Gene Repression. Cell 169, 780–791 (2017).

12. Dittmer, T.A. & Misteli, T. The lamin protein family. Genome biology 12, 222 (2011).

13. Liu, Y. et al. Tissue-specific genome editing of laminA/C in the posterior silk glands of Bombyx mori. Journal of genetics and genomics 44, 451–459 (2017).

14. Knerr, J. et al. Formin-mediated nuclear actin at androgen receptors promotes transcription. Nature 617, 616–622 (2023).

15. Kapoor, P. & Shen, X. Mechanisms of nuclear actin in chromatin-remodeling complexes. Trends in cell biology 24, 238–246 (2014).

16. Meagher, R.B., Kandasamy, M.K., Deal, R.B. & McKinney, E.C. Actin-related proteins in chromatin-level control of the cell cycle and developmental transitions. Trends in cell biology 17, 325–332 (2007).

17. Caridi, C.P. et al. Nuclear F-actin and myosins drive relocalization of heterochromatic breaks. Nature 559, 54–60 (2018).

18. Lamm, N., Rogers, S. & Cesare, A.J. Chromatin mobility and relocation in DNA repair. Trends in cell biology 31, 843–855 (2021).

19. Champion, L., Linder, M.I. & Kutay, U. Cellular Reorganization during Mitotic Entry. Trends in cell biology 27, 26–41 (2017).

20. Baarlink, C. et al. A transient pool of nuclear F-actin at mitotic exit controls chromatin organization. Nat Cell Biol 19, 1389–1399 (2017).

21. Procter, D.J., Furey, C., Garza-Gongora, A.G., Kosak, S.T. & Walsh, D. Cytoplasmic control of intranuclear polarity by human cytomegalovirus. Nature 587, 109–114 (2020).

22. Ohkawa, T. & Welch, M.D. Baculovirus Actin-Based Motility Drives Nuclear Envelope Disruption and Nuclear Egress. Current biology : CB 28, 2153–2159.e2154 (2018).

23. Zhang, X. et al. Baculovirus infection induces disruption of the nuclear lamina. Scientific reports 7, 7823 (2017).

24. Strom, A.R. et al. Phase separation drives heterochromatin domain formation. Nature 547, 241–245 (2017).

25. Larson, A.G. et al. Liquid droplet formation by HP1α suggests a role for phase separation in heterochromatin. Nature 547, 236–240 (2017).

26. Kong, X., Chen, G., Li, J., Li, Y. & Wu, X. Identification and characterization of BmNPV Bm5 protein required for the formation of nuclear vesicle structures. The Journal of general virology 104 (2023).

27. Padeken, J., Methot, S.P. & Gasser, S.M. Establishment of H3K9-methylated heterochromatin and its functions in tissue differentiation and maintenance. Nature reviews. Molecular cell biology 23, 623–640 (2022).

28. Myllys, M. et al. Herpes simplex virus 1 induces egress channels through marginalized host chromatin. Scientific reports 6, 28844 (2016).

29. Forest, T., Barnard, S. & Baines, J.D. Active intranuclear movement of herpesvirus capsids. Nat Cell Biol 7, 429–431 (2005).

30. Dai, X. & Zhou, Z.H. Structure of the herpes simplex virus 1 capsid with associated tegument protein complexes. Science 360 (2018).

31. Farrants, A. DNA Accessibility by MNase Digestions. Methods in molecular biology (Clifton, N.J.) 1689, 77–82 (2018).

32. Nagamine, T., Abe, A., Suzuki, T., Dohmae, N. & Matsumoto, S. Co-expression of four baculovirus proteins, IE1, LEF3, P143, and PP31, elicits a cellular chromatin-containing reticulate structure in the nuclei of uninfected cells. Virology 417, 188–195 (2011).

33. Nagamine, T., Kawasaki, Y. & Matsumoto, S. Induction of a subnuclear structure by the simultaneous expression of baculovirus proteins, IE1, LEF3, and P143 in the presence of hr. Virology 352, 400–407 (2006).

34. Towbin, B.D., Gonzalez-Sandoval, A. & Gasser, S.M. Mechanisms of heterochromatin subnuclear localization. Trends in biochemical sciences 38, 356–363 (2013).

35. Slack, J. & Arif, B.M. The baculoviruses occlusion-derived virus: virion structure and function. Adv Virus Res 69, 99–165 (2007).

36. Rosinski, M., Reid, S. & Nielsen, L.K. Kinetics of baculovirus replication and release using real-time quantitative polymerase chain reaction. Biotechnology and bioengineering 77, 476–480 (2002).

37. Nguyen, Q., Chan, L.C., Nielsen, L.K. & Reid, S. Genome scale analysis of differential mRNA expression of Helicoverpa zea insect cells infected with a H. armigera baculovirus. Virology 444, 158–170 (2013).

38. Shrestha, A. et al. Global Analysis of Baculovirus Autographa californica Multiple Nucleopolyhedrovirus Gene Expression in the Midgut of the Lepidopteran Host Trichoplusia ni. J Virol 92 (2018).

39. Kawasaki, Y., Matsumoto, S. & Nagamine, T. Analysis of baculovirus IE1 in living cells: dynamics and spatial relationships to viral structural proteins. The Journal of general virology 85, 3575–3583 (2004).

40. Luo, X.C. et al. Effects of Early or Overexpression of the Autographa californica Multiple Nucleopolyhedrovirus orf94 (ODV-e25) on Virus Replication. PLoS One 8, e65635 (2013).

41. Chen, L. et al. The formation of occlusion-derived virus is affected by the expression level of ODV-E25. Virus Res 173, 404–414 (2013).

42. Guo, Y.J., Fu, S.H. & Li, L.L. Autographa californica multiple nucleopolyhedrovirus ac75 is required for egress of nucleocapsids from the nucleus and formation of de novo intranuclear membrane microvesicles. PLoS One 12, e0185630 (2017).

43. Biswas, S., Willis, L.G., Fang, M., Nie, Y. & Theilmann, D.A. Autographa californica Nucleopolyhedrovirus AC141 (Exon0), a Potential E3 Ubiquitin Ligase, Interacts with Viral Ubiquitin and AC66 To Facilitate Nucleocapsid Egress. J Virol 92 (2018).

44. Mahmood, S.R. et al. β-actin dependent chromatin remodeling mediates compartment level changes in 3D genome architecture. Nature communications 12, 5240 (2021).

45. Knoll, K.R. et al. The nuclear actin-containing Arp8 module is a linker DNA sensor driving INO80 chromatin remodeling. Nat Struct Mol Biol 25, 823–832 (2018).

